# Influence of vine decline disease on the amino acid metabolism of watermelon fruit

**DOI:** 10.1101/2024.05.11.593714

**Authors:** Honoka Santo, Shota Tadano, Fumika Inokami, Takuya Nishioka, Takafumi Konaka, Motomu Sakata, Yasufumi Morimoto, Kinya Akashi

## Abstract

Vine decline (VD) is a severe disease of watermelon, melon, and other cucurbits, caused by soil-borne pathogens such as *Monosporascus cannonballus* and leads to root necrosis and sudden wilting at the later stage of fruit maturation. The present study examined the effects of VD on the metabolism of watermelon fruits. The VD-affected watermelon fruits had significantly lower lycopene and total solid contents. Still, polyphenols content and total antioxidant activities were comparable with the controls, suggesting that VD inhibited the ripening processes but maintained defensive processes in the fruits. The VD fruits showed a lower calcium level than the controls, while the contents of other major nutrition minerals were not significantly altered. The VD fruits had a lower content of total amino acids, and their composition was characterized by an increase in the percentage fractions for several amino acids, including citrulline, which may reflect the physiological response to the VD-related water deficit condition. The PCA plot clearly distinguished amino acid profiles between the VD and control fruits, demonstrating that VD exerted major influences on their amino acid metabolisms. Overall, the present study revealed that VD imposed characteristic impacts on the biochemical behaviors in the watermelon fruits.

## 1. Introduction

Watermelon (*Citrullus lanatus*) is an important vegetable fruit for the human diet. It originated from arid and semiarid regions of Africa and domesticated to the modern sweet dessert watermelon cultivars due to many years of selections (Levi et al., 2017). Watermelon accounts for 11% (100 million metric tons) of the world fruit primary production in 2022 (www.fao.org/faostat, accessed 10/5/2024), providing an important source of nutritional diets, especially in arid and semiarid regions worldwide. The high nutritional values and various health benefits of watermelon are attributable to its characteristic chemical composition, which includes a non-proteinaceous α-amino acid citrulline (Cit), lycopene, and a range of specialized metabolites (Sorokina et al., 2021).

Watermelon fruit is a rich Cit source, accumulating over 2 g per kg fresh weight in some cultivars (Hartman et al., 2019). In animals and humans, Cit can act as an arginine (Arg) precursor for the synthesis of nitric oxide, which regulates cardiovascular, immunological, and neurological functions (Bahri et al., 2013; Aguayo et al., 2021). Various health benefits of Cit consumption have been reported, which include enhancement of exercise performance and recovery (Tarazona-Díaz et al., 2013; Glenn et al., 2017), improvement of endothelial dysfunction with vasospastic angina (Morita et al., 2013), improvement of cardiac sympathovagal balance in obese postmenopausal women (Wong et al., 2016), retardation of endothelial senescence (Tsuboi et al., 2018), and alleviation of cardiovascular hypertension associated with diabetes (El-Bassossy et al., 2012). The principal natural source of Cit in the human diet is regarded as watermelon and other cucurbits since its content is either lower or undetectable in other plant/animal taxon recorded to date (Fish, 2012; Hartman et al., 2019; Aguayo et al., 2021).

In recent decades, a group of soil-borne diseases known as vine declines (VD) has been reported worldwide and caused severe production losses in cucurbits (Martyn, 1996; Rhouma et al., 2019). The major causal pathogens have been attributed to the genus *Monosporascus*, especially *M. cannonballs.* Hence the disease is commonly called *Monosporascus* root rot and vine decline (MRRVD) or *Monosporascus* sudden wilt (Martyn, 1996; Cohen et al., 2012; Rhouma et al., 2019). *M. cannonballus* appears adapted to hot, semiarid, and arid climates and grows optimally at higher temperatures over 25°C (Martyn, 1996; Cohen et al., 2012). Recent findings demonstrated that *M. cannonballus* is not the sole pathogen responsible for VD, and the disease has been associated with other root-infecting fungi such as *Monosporascus eutypoides* and *Olpidium bornovanus*, and with melon necrotic spot virus (MNSV) (de Cara et al., 2008; Stanghellini et al., 2010, 2014; Aleandri et al., 2017; Rhouma et al., 2019). This disease affects severely on melon (*Cucumis melo* L.) and watermelon, generating perithecia, which are visible as black spots on the host roots, leading to root necrosis and a sudden and severe collapse of the vines, and resulting in the wilting and death of plants in the late growing season (Martyn, 1996). The VD severely impairs fruit yield (Cohen et al., 2005; Jang et al., 2014) and fruit quality parameters such as total soluble solids and titratable acidity (Jang et al., 2014). However, the influence of VD on the chemical composition of watermelon fruit, especially on the Cit and other amino acid metabolisms, has been largely undocumented to the authors’ knowledge. Therefore, this study aimed to gain information on the amino acid metabolism of watermelon fruits affected by VD.

## 2. Materials and Methods

### 2.1. Plant materials and cultivation

Seedlings of watermelon (*Citrullus lanatus* L.) cultivar Matsuribayashi RG (Hagihara Farm, Tawaramoto, Nara, Japan), which was grafted to a wild-type watermelon (*Citrullus lanatus* sp.) rootstock cultivar Donna-Mondai (Tottori Prefectural Horticultural Research Center, Hokuei, Tottori, Japan), were used in this study. The grafted seedlings in pots were transplanted on March 21, 2017 on the ground, in two plots at least 8 m apart, with clay loam soil in polyolefin greenhouse facilities at the Tottori Prefectural Horticultural Research Center, with 0.5 m spacing within rows and 3.0 m between rows. A compost at a rate of 3 kg m^−2^, and chemical fertilizers 140 g m^−2^ of Seruka-Friend (Zen-Noh, Tokyo, Japan) and 90 g m^−2^ of Suika-Ippatsu (Zen-Noh) were applied before planting, which corresponded to the application of N: P_2_O_5_: K_2_O dose of 12.6:12.0:2.7 g m^−2^. Standard practice for cultivation management was applied. A spontaneous and unintentional VD, diagnosed by a sudden and severe wilting of their vines, occurred accidentally in one of the two plots at a later ripening stage. Watermelon fruits in the VD-affected plot were nevertheless harvested on June 13–15, 2017, approximately 50 days after pollination, and used as diseased samples, while fruits from asymptomatic, morphologically healthy watermelon individuals from the other plot were used as controls. The transverse and longitudinal diameters of the fruits were measured using a ruler, and the fruit was weighed using an electric balance.

### 2.2. Soluble solid content and flesh sample preparation

Fruits were cut in half, and soluble solids content (SSC) of the flesh at the central position of the fruit and in the vicinity of seeds was immediately measured with a digital refractometer (PR101α, ATAGO, Tokyo, Japan) as degree Brix (°Bx) value. For other biochemical analyses, flesh at the central position of the fruit was cut into a cube of 20 mm side and stored in a −30°C freezer until extractions. The cubes were homogenized using a mortar and pestle in the presence of liquid nitrogen. Then the homogenates were dispensed into 200 μl aliquots in 1.5 ml plastic tubes and stored in a −30°C freezer until further chemical analyses.

### 2.3. Lycopene assay

Lycopene assay was performed as described previously (Akashi et al., 2017) with the following modification. The 100 mg of the homogenate was mixed with 8 ml of HAE-BHT solution (hexane:acetone:ethanol, (2:1:1(v/v)), supplemented with 0.05% butylated hydroxytoluene) in a 15 ml plastic tube. Three 5 mm- and one 10 mm stainless steel beads were added. Then, the tube was placed horizontally in a shaking incubator (MildMixer PR-36, Taitec, Koshigaya, Saitama, Japan) and shaken at medium speed for 10 minutes at 25°C. Then, 1.2 mL of water was added and shaken in the shaking incubator for 10 min. The homogenate was centrifuged at 8,000 *g* for 10 min at 25°C, and the optical absorbance of the supernatant was measured at 503 nm wavelength in a spectrophotometer (UH5300, Hitachi High-Tech, Tokyo, Japan). Lycopene content was calculated using the molecular absorption coefficient of lycopene (172,000 M cm^−1^) (Perkins-Veazie et al., 2001).

### 2.4. Total phenols content and antioxidant activity

The 165 mg of the watermelon flesh homogenate was placed into a 1.5 mL tube, 385 µl of ethanol was added and vortexed vigorously for 1 h using a shaker (MT-400, Tomy, Digital Biology, Tokyo, Japan). The 300 µl of the mixture was transferred to a new 1.5 ml tube, and 300 µl of 70% ethanol was added and vigorously vortexed. The mixture was centrifuged at 20,000 *g* for 2 min at 4°C, and the supernatant was used as an extract for total polyphenols content and antioxidant activity assays. Total polyphenol content was measured as described previously (Maalej et al., 2017) with the following modifications. The watermelon flesh homogenate (40 µl) or standards (gallic acid in the range of 0–80 µg ml^−1^) were mixed with 200 µl of 10% Folin-Ciocalteu’s phenol reagent (Sigma-Aldrich, Merck, Burlington MA, USA) and incubated at 25°C in the dark for 5 min. The 160 µl of 7.5% sodium carbonate in H_2_O was added and incubated at 25°C for 1 hr in the dark. The 300 µl of the reactant was transferred to a 96-well plate, and the absorbance at 750 nm was measured using a plate reader (Multiskan FC, ThermoFisher, Waltham, MA, USA). The total polyphenol content was expressed as gallic acid equivalent per gram of watermelon flesh sample.

The antioxidant activity of the extract was assayed as 1,1-phenyl-2-picrylhydrazyl (DPPH) radical scavenging activity essentially as described previously (Maalej et al., 2017) with the following modifications. The watermelon flesh homogenate (200 µl) or standards (Trolox (Tokyo Chemical Industry, Tokyo, Japan) in the range of 0–80 nmol) were mixed with 200 µl of 300 µM DPPH (Tokyo Chemical Industry) solution in ethanol, and incubated at 25°C in the dark for 1 hr. The 300 µl of the reactant was transferred to a 96-well plate, and the absorbance at 520 nm was measured using the Multiskan plate reader. Antioxidant activity was expressed as Trolox equivalent per gram of watermelon flesh sample.

### 2.5. Mineral contents

Mineral contents of the watermelon flesh sample were determined as described previously (Yamada et al., 2018) with the following modifications. The homogenized watermelon flesh sample (0.5 g) was placed into a 50 ml Erlenmeyer flask, to which 10 ml of nitric acid (Electric industry grade, 70%, density 1.42 g ml^−1^, Fujifilm Wako Pure Chemical, Osaka, Japan) was added. The mixture was digested on a hot plate for 1 h each at 90, 140, 190, and 240°C, and then the digested sample was diluted to 25 ml with 1% nitric acid. The amount of mineral elements was determined using an inductively coupled plasma atomic emission spectroscopy (Spectro Ciros CCD, Spectro Analytical Instruments, Nordrhein-Westfalen, Germany), using the Metal Standard Solutions (registered by the Japan Calibration Service System, Fujifilm Wako Pure Chemical) as reference materials.

### 2.6. Amino acid analysis

Extraction of amino acids from watermelon samples was performed as described previously (Kawasaki et al., 2000) with the following modifications. The 100 mg of homogenized watermelon flesh was mixed with 650 µl of extraction buffer (methanol:chloroform:water (5:2:2 (v/v))) and vortexed for 5 min. The extract was centrifuged at 20,000 *g* for 2 min at 4°C, and the supernatant was transferred into a new tube. The 195 µl of chloroform and 195 µl of water were added and vortexed for 5 seconds. The extract was centrifuged again at 20,000 *g* for 2 min at 4°C, and the supernatant was filtered through a hydrophilic polytetrafluoroethylene membrane filter (Millex-LG, 0.2 um × 4 mm, Merck Millipore, Darmstadt, Germany) and transferred to a screw-top glass vial. Amino acid contents were measured using a triple quadrupole LC-MS/MS system (Agilent 6420, CA, USA) with a Discovery HS-F5 column (2.1 × 250 mm, 5 µm, Sigma-Aldrich) as described previously (Itam et al., 2020), with a mobile-phase gradient generated by 0.1% formic acid and acetonitrile as detailed in Supplementary Document S1. The 33 metabolites were used as standards and dissolved in 0.1% formic acid in either water or 50% methanol to make a 10 ppm amino acid mixture. Metabolites were identified by multiple reaction monitoring analysis, with characteristic product ions as shown in Supplementary Table S1.

### 2.7. Statistical analyses

In each assay, each fruit specimen was derived from an independent plant individual, thus statistical validations in this study were based on the biological replicates. The Student’s *t*-test was performed using the Microsoft Excel software (Redmond, WA, USA). One-way ANOVA with post-hoc Tukey HSD testing was performed using the Astatsa.com online statistical calculator (Astatsa, 2016). Principal component analysis (PCA) was performed using the prcomp function in the stats package (version 3.6.2) in R statistical software (R Core Team, 2016).

## 3. Results

### 3.1. Effect of VD on fruit size, weight, and total solid content

In watermelon cultivation in 2017, an accidental VD disease outbreak occurred spontaneously in one of the two plots at the Tottori Prefectural Horticultural Research Center. The symptoms were typical of MRRVD, with a sudden wilting of the leaves, collapse of the vines, and reduction of fruit load (Martyn, 1996; Cohen et al., 2012). Black spots, characteristic of MRRVD, were observed on the roots of the affected watermelon (Martyn, 1996). Watermelon plants in the other plot were morphologically normal, and no disease symptoms were observed. Transverse and longitudinal diameters of fruits harvested from VD-affected watermelon plants were 24.2 ± 0.5 and 24.6 ± 1.3 cm, respectively, in contrast to those from the asymptomatic healthy watermelon plants (26.6 ± 0.5 and 29.7 ± 1.1 cm, respectively), which resulted in the 8.9 and 17.1% reduction, respectively (Fig. 1a–b). Fresh weight of the fruits also showed a decline of 32.7% in VD-affected watermelon (11.1 ± 1.0 and 7.5 ± 0.6 for control and VD-affected fruits, respectively, Fig. 1c).

**Figure 1.**
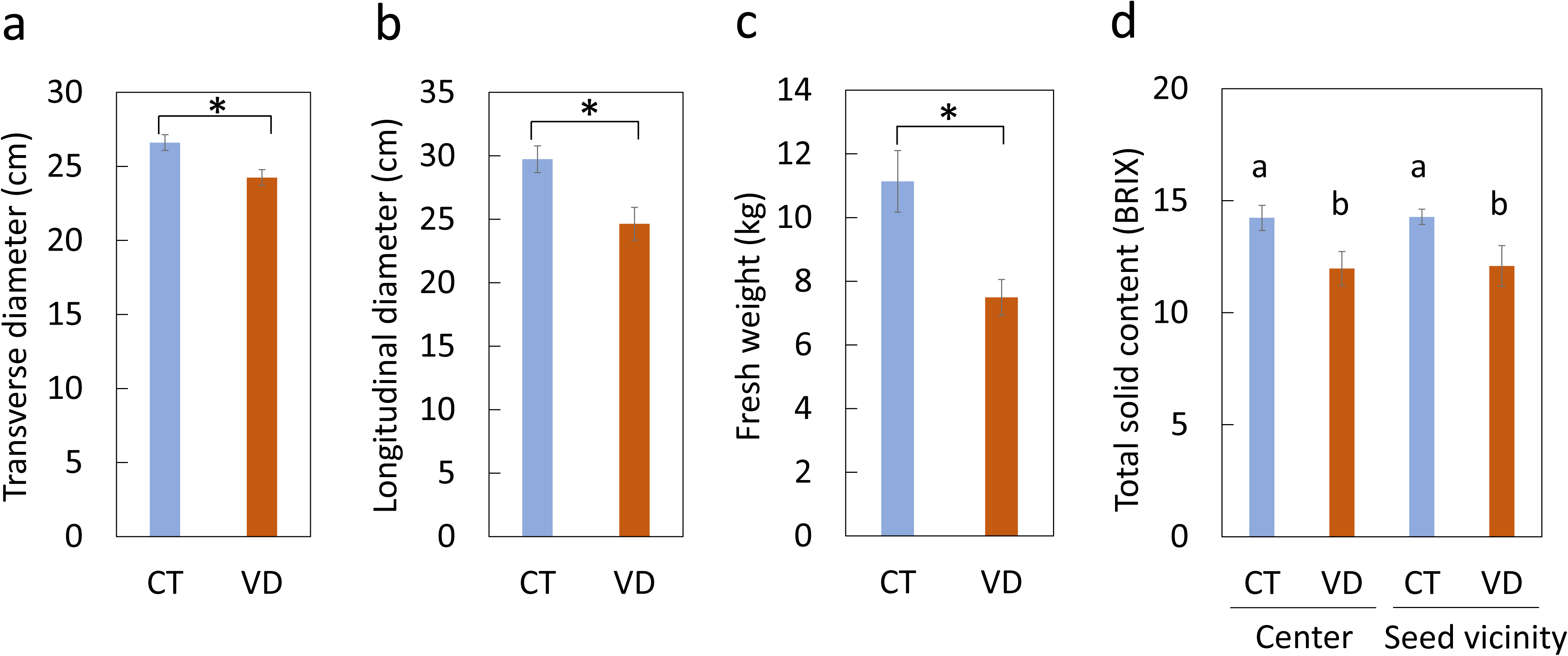
Effect of VD on transverse (a) and longitudinal (b) diameters, fresh weight(c), and soluble solid content (d) in watermelon fruits. CT and VD represent fruits from non-symptomatic control and VD-affected watermelon plants. To analyze total solid contents, watermelon flesh at the central position of flesh and in the vicinity of seeds were measured. Data represent means and standard deviations for n=7–9 from independent plant individuals. In (a–c), statistically significant differences at a 5% level by t-test are indicated by asterisks. In (d), statistically different values are indicated by different alphabets using one-way ANOVA with post-hoc Tukey HSD test (p<0.05).

The total solid content of the flesh as measured by the BRIX value, which reflects sugar accumulation in the fruit (J. Huang et al., 2022), dropped by 15.9 and 15.4% at the central and seed vicinity position, respectively, in the flesh of the VD-affected fruit in comparison to the healthy controls (Fig. 1d).,

### 3.2. Lycopene and polyphenol contents and antioxidant activity

Lycopene is a carotenoid with significant antioxidant activity and has gained attention for its benefits to human health (Grabowska et al., 2019; Sorokina et al., 2021). The lycopene content in the central position of flesh decreased by 52% in the VD-affected watermelon (42.7 ± 3.3 and 20.7 ± 4.1 μg gFW^−1^, respectively, Fig. 2a). By contrast, content of polyphenols, a group of antioxidative plant specialized compounds with multiple health benefits (Abbas et al., 2017; Luca et al., 2020), was statistically not different to those in the control watermelon at the 5% level (59.7 ± 8.5 and 54.7 ± 7.9 μg gFW^−1^ for the control and VD-affected watermelon, respectively, *P*=0.135 by *t*-test, n = 7–9, Fig. 2b). Total antioxidant activity, as examined by the DPPH free radical quenching assay (Sharma & Bhat, 2009), was also statistically equivalent to those in the control watermelon (71.5 ± 9.4 and 63.9 ± 14.5 µmol Trolox equivalent gFW^−1^ for control and VD-affected plants, respectively, *P*=0.202 by *t*-test, n = 5, Fig. 2c).,

**Figure 2.**
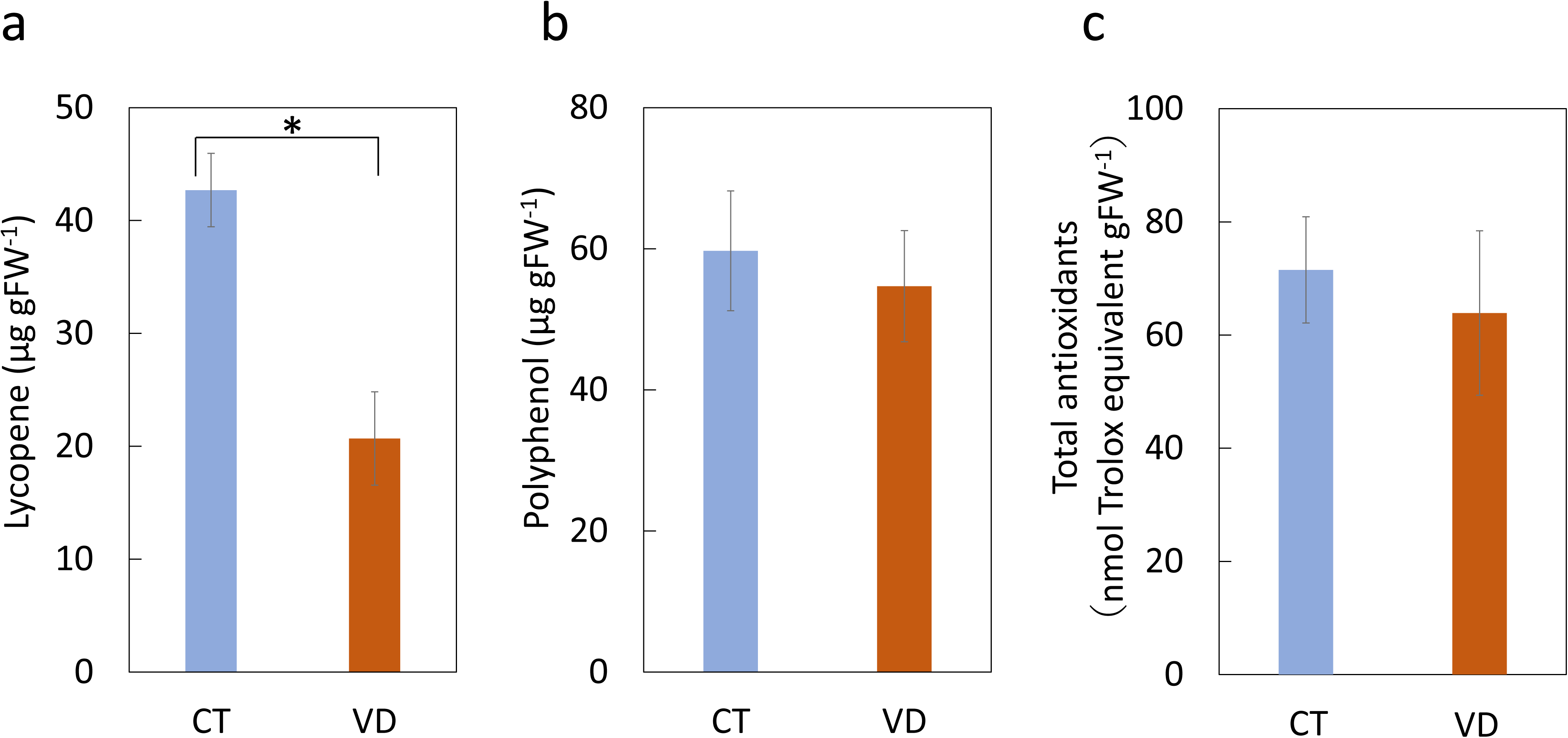
Contents of lycopene (a), polyphenols (b), and total antioxidants (c) in the central position of watermelon flesh harvested from non-symptomatic control plants (CT) and VD-affected plants (VD). Data for lycopene contents represents means and standard deviation for 6 independent plant individuals, while those for polyphenols and total antioxidants were obtained from 7–9 independent plant individuals. Statistically significant differences at the 5% level by *t*-test are indicated by asterisk.

### 3.3. Mineral content

Since VD primarily impairs root function in the infected watermelon plants (Martyn, 1996; Cohen et al., 2012; Rhouma et al., 2019), absorption and accumulation of plant nutrition minerals in the fruits may be impaired in the VD-infected watermelon. The contents of major nutrient minerals in the fruits were analyzed to address this possibility. Watermelon fruit has been known for its high potassium (K) content (Fulgoni & Fulgoni, 2022). In the present study, K content in the MRRVD-affected fruits was not statistically different from the control at the 5% level (996 ± 231 and 675 ± 208 ng gFW^−1^ for the control and VD-affected fruits, respectively, *P* = 0.098 by *t*-test, n = 4–5, Fig. 3a). Similar trends were observed for most of the other minerals (Fig. 3), except for calcium (Ca), which showed statistically significant decrease by 31% in the VD-affected watermelon in comparison to the control (48.6 ± 7.9 and 33.4 ± 8.9 ng gFW^−1^ for the control and VD-affected fruits, respectively).,

**Figure 3.**
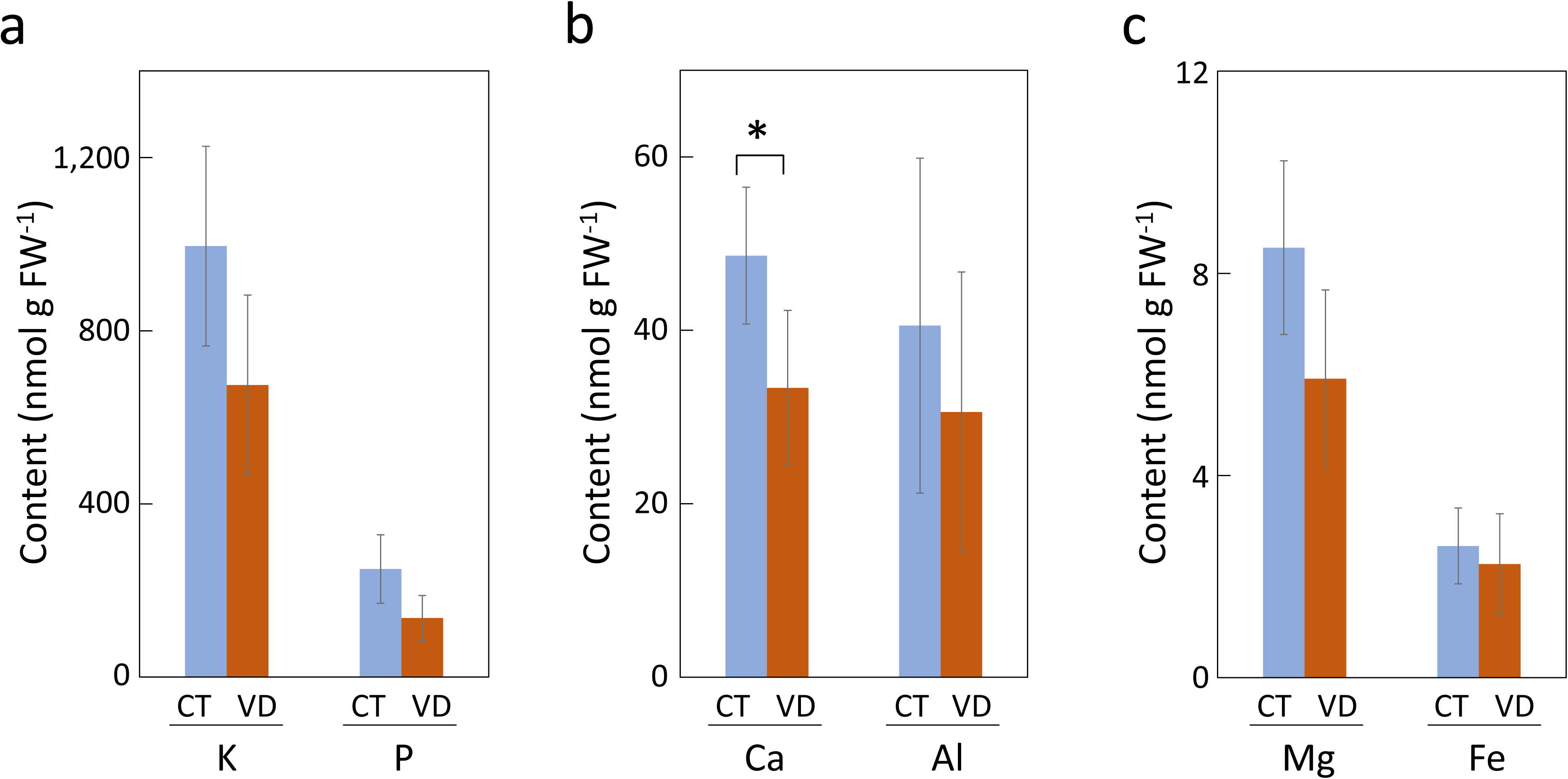
Contents of major minerals in the center of watermelon fruit flesh harvested from control watermelon plants (CT) and VD-affected (VD) watermelon plants. Data represent means and standard deviations obtained from 4–5 independent plant individuals. Data are the contents for potassium (K), phosphorus (P) (a), calcium (Ca), aluminum (Al) (b), magnesium (Mg), and iron (Fe) (c). Statistically significant differences at the 5% level by *t*-test are indicated by asterisks.

### 3.4. Free amino acid composition

LC-MS/MS analysis detected 22 free amino acids from the VD-affected and control watermelon fruits (Fig. 4a–c, Supplementary Table S2). Total free amino acid content was decreased from 26.03 ± 1.67 mg gFW^−1^ in the control to 20.29 ± 2.60 mg gFW^−1^ in the VD fruit, resulting in a 22.0% decline under the disease condition (Supplementary Table S2). Cit accumulated 7.08 ± 0.61 and 8.27 ± 1.13 mg gFW^−1^ in the control and VD-affected fruits, respectively, occupying the largest percentage fractions of 27.19 ± 1.54 and 38.14 ± 2.67%, respectively under control and VD conditions (Fig. 4d–e, Supplementary Tables S2–S3). Although the Cit content as a fresh weight basis was not statistically different between the VD and control conditions at the 5% level (*P* = 0.17 by *t*-test, n = 5, Fig. 4a), its percentage fraction significantly increased in the VD watermelon than in the control (Supplementary Table S2).

**Figure 4.**
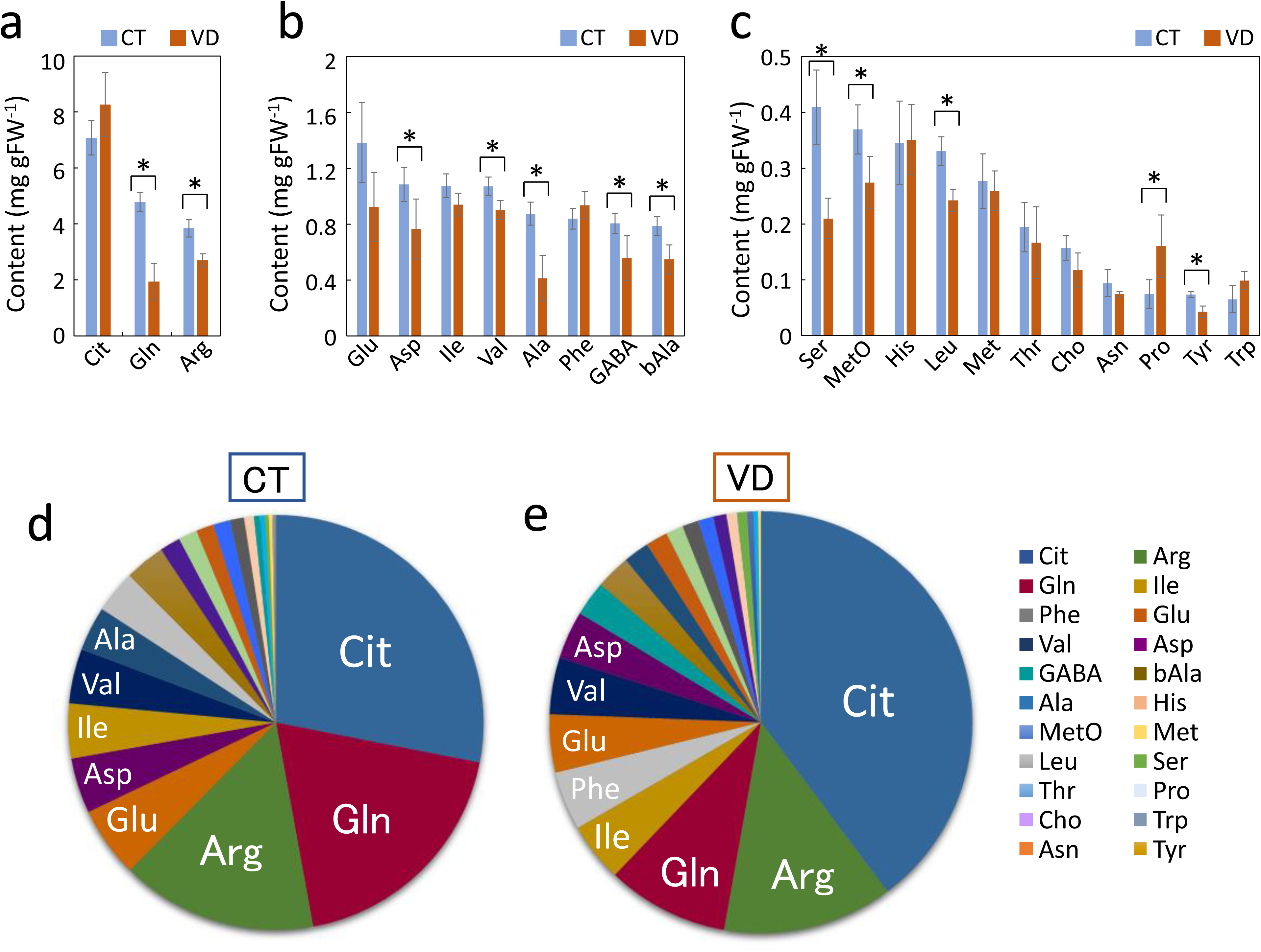
Amino acid content and composition in the center of watermelon flesh. The amino acids in the range over 2 mg gFW^−1^ (a), 0.5–2 mg gFW^−1^ range, and below 0.5 mg gFW^−1^ (c) are shown. Data represent means and standard deviations for 5 independent plant individuals for control (CT) and VD-affected (VD) fruits. In (a)–(c), statistically significant differences between CT and VD at the 5% level by *t*-test are indicated by asterisks. Pie chart representation for the amino acid composition for the control (d) and VD-affected fruits (e). Data are the average for n=5 for each chart.

By contrast, contents of glutamine (Gln) and Arg, which occupied the second and third largest concentration in control watermelon (4.79 ± 0.34 and 3.85 ± 0.31 mg gFW^−1^ for glutamine and arginine, respectively), decreased significantly in the VD-affected watermelon (1.94 ± 0.65 and 2.70 ± 0.24 mg gFW^−1^, respectively). A significant decrease in the accumulation level was observed in many of the other amino acids such as aspartate (Asp), valine (Val), alanine (Ala), γ-aminobutyrate (GABA), beta-alanine (bAla), serine (Ser), methionine sulfoxide (MetO), leucine (Leu), and tyrosine (Tyr) (Fig. 4b–c). Proline (Pro) was unique in that its accumulation level at the fresh weight basis was significantly elevated by 2.16-fold in the VD-affected watermelon (0.074 ± 0.025 and 0.160 ± 0.056 mg gFW^−1^ in the control and VD-affected fruits, respectively, Fig. 4c and Supplementary Table S2). However, the percent fraction was only 0.25 and 0.5% of the total amino acids in the control and VD fruits, respectively (Supplementary Table S3).

The percentage fraction of each amino acid and its pie chart showed a significant increase in Cit, isoleucine (Ile), phenylalanine (Phe), histidine (His), Pro, and tryptophan (Trp) in the VD fruits (Fig. 4d–e and Supplementary Table S3), while decreased percentage fractions in the VD in comparison to the control were observed for Gln, Ser, and Tyr.

A metabolic pathway diagram was illustrated to visualize the behaviors of amino acid accumulation in the VD-affected watermelon (Fig. 5). The diagram indicated that the amino acids in the same and/or closely related pathways often show contrasting accumulation behaviors. For example, Cit showed an increased percentage fraction, while its downstream metabolite Arg exhibited a decrease in concentration in the VD watermelon. In another case, Trp and Phe in the aromatic amino acids (AAAs) pathway showed an increase in percentage fractions, while Tyr in the same pathway exhibited a decrease in both concentration and percentage fraction in the VD fruit.

**Figure 5.**
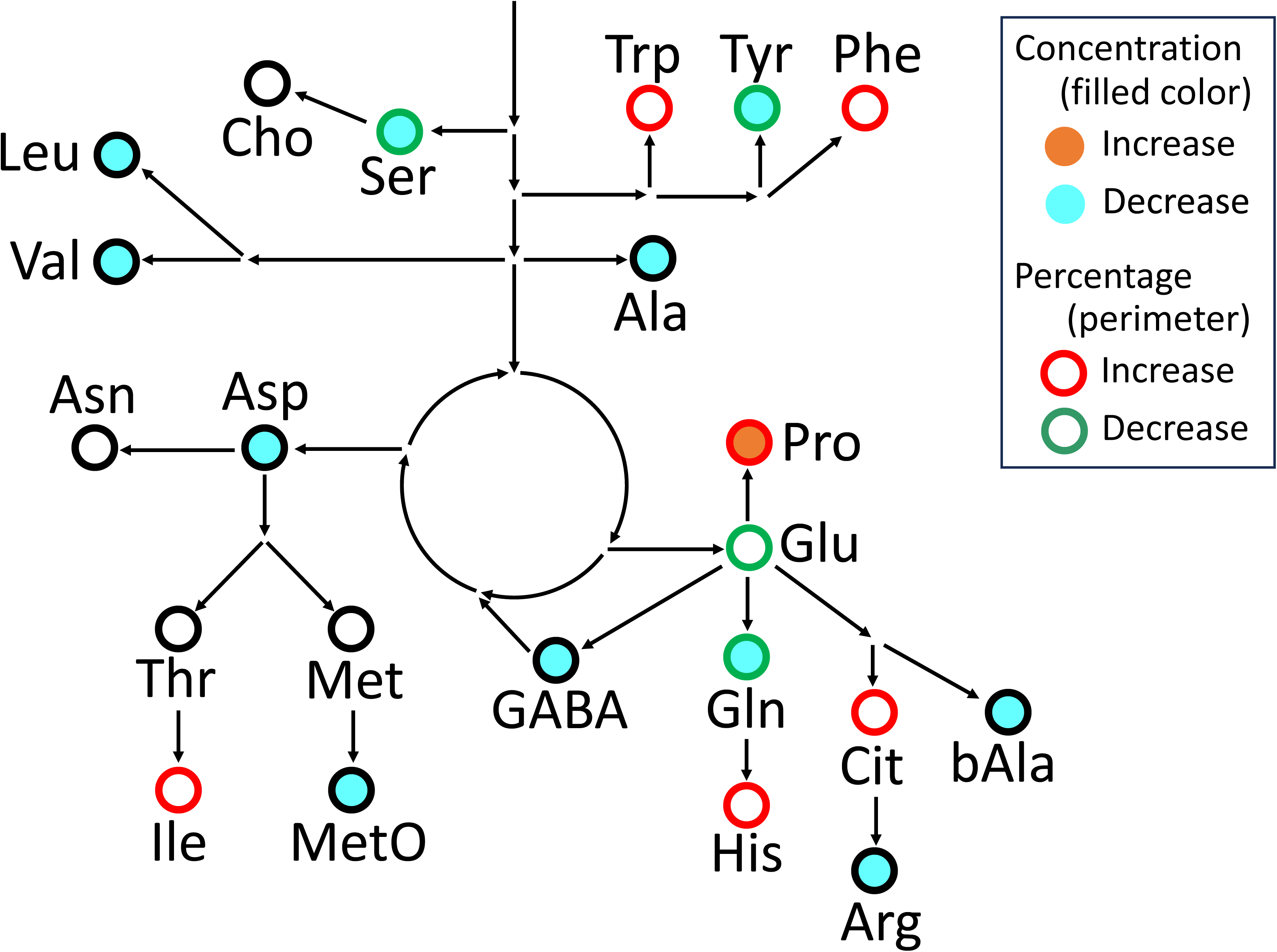
A schematic view of the metabolic pathway highlights the change in free amino acid accumulation levels in the VD-affected fruits. Amino acids, which significantly increased and decreased concentration at the fresh weight basis in the VD-affected watermelon fruits (P<0.05 by t-test), are shown in orange and sky blue circles, respectively. Amino acids within which the percent fraction increased and decreased (P<0.05 by *t*-test) in the VD watermelon are shown in the red and green perimeter of the circles, respectively. Amino acids with a black perimeter of the circles indicate that their difference in the percent fraction between VD and control watermelon was not statistically significant at the 5% level.

### 3.5. Principal component analysis (PCA)

The free amino acids profile in the VD-affected and the control watermelon fruits were further analyzed by PCA. Consequently, clear separation of VD-affected and control watermelon fruits was observed in the score plot, in which VD-affected fruits were located on the PC1-negative side, mainly in the quadrant III location. In contrast, control fruits were situated on the PC1-positive side (Figure 6). A loading plot of the PCA suggested that several amino acids, notably Trp, Pro, and Phe, were associated with VD, while most of the other amino acids tended to cluster with control fruits.

**Figure 6.**
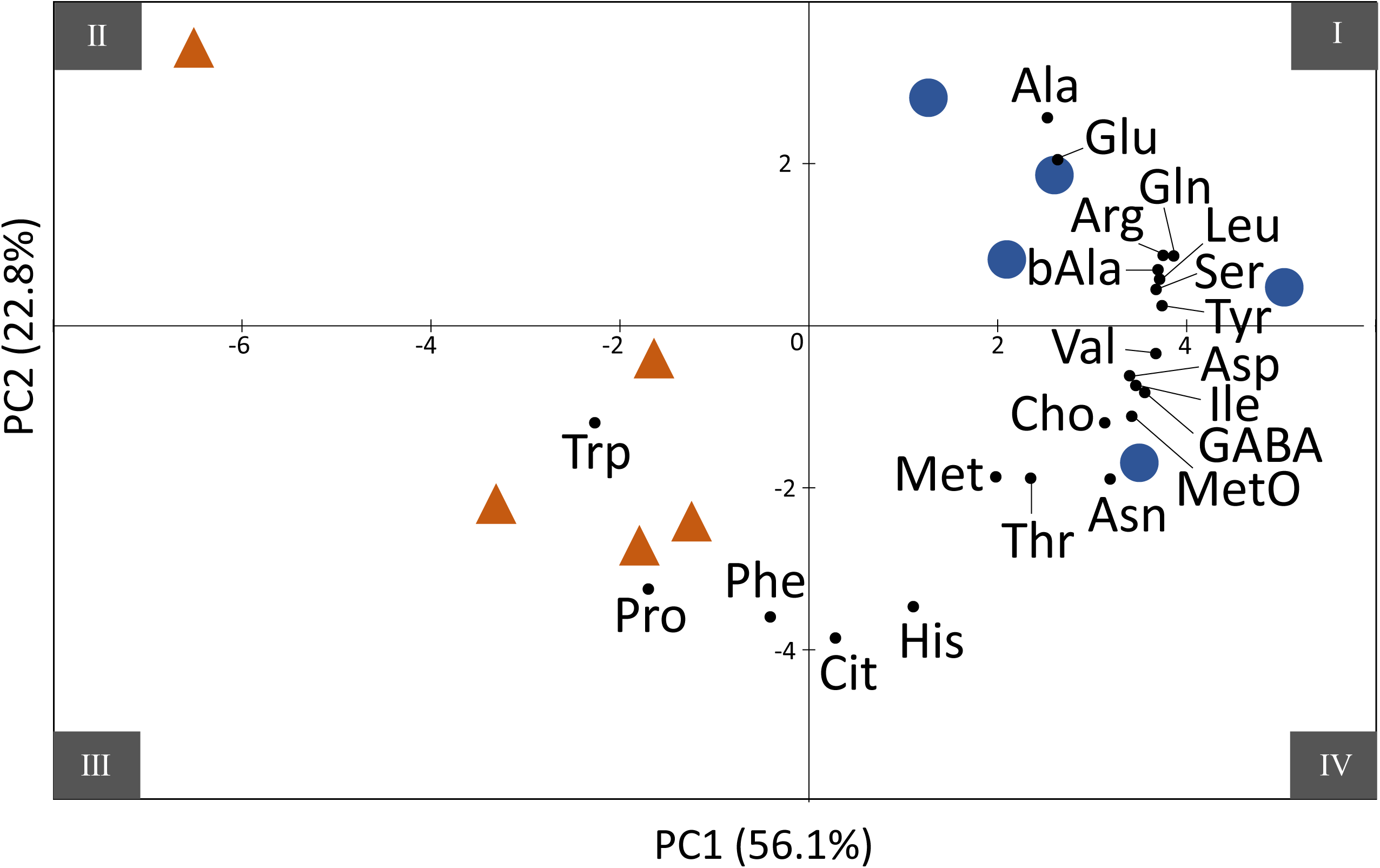
Principal component analysis of free amino acid accumulation patterns in the VD-affected and control watermelon fruits. Scores for the VD-affected (orange triangles) and control (blue circles) fruits from individual watermelon plants (n=5) are plotted in the PC1-PC2 space. Locations for the quadrants I to IV are indicated at the four corners of the plot. Small black dots show directions and relative strength of the loading values for the respective amino acids.

## 4. Discussion

A disease that causes sudden wilting or VD symptoms, commonly known as MRRVD, is recognized as one of the most important diseases in watermelon and melon and is a limiting factor in their production in some of the cultivation zones around the world (Cohen et al., 2012; Rhouma et al., 2019). While many reports have been documented on the pathogenic actions and yield impact of VD, less has been reported on the impact on fruit metabolism and quality. The present study showed that VD influenced the composition of specific metabolites in the affected fruits.

The fruits affected with VD showed a different free amino acid profile from the control watermelon (Fig. 4–6, Supplementary Table S2–S3). The percentage fraction of Cit, an amino acid predominant in watermelon fruits (Hartman et al., 2019), was shown to be increased, while Gln and Arg, the second and third largest constituents in the fruit, decreased in the VD-affected fruits. This behavior is reminiscent of previous reports on amino acid metabolism in watermelon vegetative tissues under drought stress, in which the elevation of Cit level was regulated at the transcriptional and enzyme abundance of the Cit/Arg pathway (Kawasaki et al., 2000; Song et al., 2020). Present observations were consistent with previous reports in which xylem hydraulic conductance was decreased in MRRVD-affected melon plants via tyloses development in roots, causing wilting symptoms similar to drought stress (Pivonia et al., 2002; Cohen et al., 2012). In plants, Cit is implicated in nitrogen storage and transport (Ludwig, 1993) and the scavenging hydroxy radical (Akashi et al., 2001), a toxic reactive oxygen species under stress conditions. Therefore, maintenance of the high level of Cit in the VD-affected watermelon may suggest the presence of a common defense/adaptive mechanism against water-deficit physiological conditions under both abiotic and biotic stresses.

Notably, Pro was the only amino acid that increased in abundance at the fresh weight basis in the VD fruits (Fig. 4c and Supplementary Table S2). Increased accumulation of Pro in response to various environmental stresses is widely observed in diverse plant species (Verbruggen & Hermans, 2008; Ghosh et al., 2022). Pro has long been known as an osmoprotectant under water-deficit conditions in plants (Mattioli et al., 2009; Ghosh et al., 2022). In this study, however, the Pro accumulation level in VD-affected watermelon fruits observed was only 0.160 ± 0.056 mg gFW^−1^, far lower than the 1–8 mg gFW^−1^ accumulation ranges for other major amino acids Cit, Gln, and Arg (Fig. 4 and Supplementary Table S2), suggesting that the induced Pro is not likely to contribute as an osmoprotectant in the fruit. Recently, multifaceted functions of Pro have been proposed, such as a chelator for heavy metals, a molecular chaperone, a reactive oxygen species scavenger, a regulator of redox balance and energy status, and a signal messenger (Szabados & Savouré, 2010; Trovato et al., 2021; Zheng et al., 2021; Ghosh et al., 2022). The induced Pro in the VD fruits may also have a non-osmoprotectant physiological significance, although its elucidation awaits further investigation.

The present study revealed contrasting behaviors of AAAs levels in the VD-affected watermelon fruit, where Tyr content decreased, while the percent fraction of Phe and Trp increased in the affected fruit (Fig. 4c and 5, Supplementary Table S2–3). The physiological significance of the different behaviors of these amino acids is currently unknown. Still, one scenario is that changes in their accumulation ratios may reflect their differential requirements as precursors for downstream metabolisms. AAAs serve as precursors for protein synthesis and numerous plant-specialized metabolites in plants (Maeda & Dudareva, 2012; Yokoyama et al., 2021; Almeida et al., 2023). In watermelon, many alkaloids, shikimates, phenylpropanoids, and polyphenols are detected (Sorokina et al., 2021). For example, luteolin, chlorogenic acid, rutin, quercetin, vanillic and sinapic acids are reported as the main phenolics found in watermelon fruits, which are synthesized from Phe (Fu et al., 2011; Mushtaq et al., 2015; Abu-Hiamed, 2017; Haytovitz et al., 2018; Ilahy et al., 2019). Watermelon fruits contain Trp-derived alkaloids such as melatonin and serotonin (Islam et al., 2016; Mandal et al., 2018), which function in stress signal transduction and mitigation of abiotic/biotic stresses including pathogen attack (Wang et al., 2020; Chang et al., 2021; Wu et al., 2021). Unraveling the relationships between these plant-specialized metabolites and their AAAs precursors in the VD-affected fruits should await further investigation.

This study also showed that lycopene and total solid contents were markedly lower. In contrast, polyphenol content was unchanged in the VD-affected watermelon fruits compared to the controls (Fig. 2a–b). Lycopene is one of the natural tetraterpenoid pigment family carotenoids, which play diverse and essential functions in plants, such as photosynthesis, photoprotection, biosynthetic precursors for phytohormones, and signal transduction (Sun et al., 2018). Pigmentation of flowers, fruits, and seeds by carotenoids is also an important trait to attract pollinators and seed dispensers, thereby facilitating successful reproductive processes in plants (Valenta et al., 2018; Sun et al., 2022). Lycopene represents a major carotenoid in red-fleshed watermelon fruits, accumulating at the fruit-loading stage via transcriptional and post-transcriptional regulations (Zhang et al., 2020; Yuan et al., 2021). Soluble sugars such as sucrose and fructose also accumulate at the loading stage in watermelon fruits (Liu et al., 2013; Ren et al., 2023). The decrease in lycopene and total solid contents in the VD-affected watermelon fruits is consistent with previous observations that MRRVD inhibits the ripening processes of watermelon fruits (Martyn, 1996; Cohen et al., 2012). By contrast, the unchanged content of polyphenols suggested that their accumulations may precede the growth perturbation by VD, and/or their de novo synthesis may be maintained under the VD-induced growth perturbation. Polyphenols have potent antioxidative activities and play a role in the defense processes against abiotic/biotic stresses (Abbas et al., 2017; Luca et al., 2020), which may explain the sustained level of total antioxidants in the VD-affected watermelon (Fig. 2c).

Effects of VD on the accumulation levels of nutrition minerals were not obvious in the present study (Fig. 3). Although decreases in the mean values for six major minerals were observed in the VD-affected fruits compared to the controls, the values were highly variable between fruits. Their differences were not statistically validated except for Ca, which decreased by 31% in the VD-affected fruits (Fig. 3b). In plant fruits, Ca is implicated in diverse functions such as signal transduction (Fenn & Giovannoni, 2021), and cross-linking-mediated structural reinforcement of the cell walls (Chan et al., 2017; W. Huang et al., 2023; Winkler et al., 2020), raising a possibility that the observed Ca decline may adversely affect ripening process in the VD-affected fruits. Mineral metabolism and its physiological implications under VD should be further addressed in future studies.

In this study, we examined fruits from the VD-affected watermelon plants, which occurred spontaneously and unintentionally in one of the two experimental plots in the research station. The observed morphological symptoms were typical for MRRVD, characterized by a sudden and severe wilting of their vines at a later ripening stage and black spots on the roots. However, this study has not done anatomical and/or genetic verifications for MRRVD; thus, further studies are required to determine the effect of MRRVD on watermelon fruit metabolism. Moreover, the possibility that differences in the soil microenvironments between the two plots may have affected the observed biochemical characteristics of the fruits cannot be excluded. To address these issues, further experiments with controlled MRRVD infection in an appropriate field block design will be needed in future studies.

## 5. Conclusions

VD, caused by soil-borne pathogens such as *M. cannonballus,* has a serious impact on watermelon cultivation and production around the world. Much research has been reported on VD’s pathological actions, disease control cultivation management, and disease-resistant varieties/seedlings establishment (Martyn, 1996; Cohen et al., 2005, 2012). However, few studies have been reported on the biochemical aspects of how VD affects the nutritional composition of watermelon fruits. The present study revealed that VD has a distinctive effect on the chemical composition of watermelon fruit flesh. Decreased total solid contents and lycopene were observed in the fruits affected by VD, suggesting that the disease-induced growth impairment inhibited fruit maturation. However, polyphenol levels and total antioxidant activity were maintained in the VD-affected fruits, which may indicate the activation and/or maintenance of defense mechanisms against pathogen attacks. The Ca content declined in the affected fruits, which may influence Ca-related cellular functions in the VD-affected fruits. Although the level of total free amino acids was decreased, the increase in the percent fraction of Cit was observed in the VD-affected fruits, which may be relevant to the response against water deficit and wilting caused by the disease. Contrasting behaviors of AAAs were observed in the VD-affected fruits, characterized by the decrease in Tyr content and the increased percentage fractions of Phe and Trp, implying the differential demands and metabolic regulations thereof for these amino acids under disease-affected conditions. The PCA plot showed a clear distinction of amino acid profiles between the VD and control fruits, demonstrating that VD exerted major influences on the amino acid metabolisms in fruits. Overall, the present study revealed that VD imposed characteristic impacts on the biochemical behaviors in the watermelon fruits, which offers a new insight into the mode of action of VD in the cucurbits.

## Author Contributions

Conceptualization: HS, YM, and KA. Data curation: HS, YM, and KA. Formal analysis: HS, ST, and KA. Funding acquisition: YM and KA. Investigation: HS, YM, and KA. Methodology: HS, ST, FI, TN, TK, MS, and KA. Project administration: HS, YM, and KA. Resource: YM. Software: HS, ST, and KA. Supervision: YM and KA. Validation: KA. Visualization: HS and KA. Writing – original draft: HS. Writing – review and editing: ST, FI, TN, TK, MS, YM, and KA.

## Data availability statement

The data supporting this study’s findings are available upon request from the corresponding author, [K. A.].

## Acknowledgments

We thank the staff members of the Laboratory of Molecular and Cellular Biology, Faculty of Agriculture, Tottori University, and of the Tottori Prefectural Horticultural Research Center, Tottori Prefecture, for technical support in the laboratory and in the field.

## References

Abbas, M., Saeed, F., Anjum, F. M., Afzaal, M., Tufail, T., Bashir Muhammad Shakeel, Ishtiaq, A., Hussain, S., & Suleria, H. A. R. (2017). Natural polyphenols: An overview. 10.1080/10942912.2016.1220393

Abu-Hiamed, H. (2017). Chemical composition, flavonoids and β-sitosterol contents of pulp and rind of watermelon (*Citrullus lanatus*) fruit. Pakistan Journal of Nutrition, 16, 502–507. 10.3923/pjn.2017.502.507

Aguayo, E., Martínez-Sánchez, A., Fernández-Lobato, B., & Alacid, F. (2021). L-Citrulline: A non-essential amino acid with important roles in human health. Applied Sciences, 11(7), 3293. 10.3390/app11073293

Akashi, K., Mifune, Y., Morita, K., Ishitsuka, S., Tsujimoto, H., & Ishihara, T. (2017). Spatial accumulation pattern of citrulline and other nutrients in immature and mature watermelon fruits. Journal of the Science of Food and Agriculture, 97(2), 479–487. 10.1002/jsfa.7749

Akashi, K., Miyake, C., & Yokota, A. (2001). Citrulline, a novel compatible solute in drought-tolerant wild watermelon leaves, is an efficient hydroxyl radical scavenger. FEBS Letters, 508(3), 438–442. 10.1016/S0014-5793(01)03123-4

Aleandri, M. P., Martignoni, D., Reda, R., Alfaro-Fernández, A., Font, M. I., Armengol, J., & Chilosi, G. (2017). Involvement of *Olpidium Bornovanus* and *O. Virulentus* in the occurrence of melon root rot and vine decline caused by *Monosporascus Cannonballus* in central Italy. Journal of Plant Pathology, 99(1), 169–176. 10.4454/jpp.v99i1.3787

Almeida, A. M., Marchiosi, R., Abrahão, J., Constantin, R. P., dos Santos, W. D., & Ferrarese-Filho, O. (2023). Revisiting the shikimate pathway and highlighting their enzyme inhibitors. Phytochemistry Reviews. 10.1007/s11101-023-09889-6

Astatsa. (2016). *Online web statistical calculators* [Computer software]. https://astatsa.com/

Bahri, S., Zerrouk, N., Aussel, C., Moinard, C., Crenn, P., Curis, E., Chaumeil, J.-C., Cynober, L., & Sfar, S. (2013). Citrulline: From metabolism to therapeutic use. Nutrition, 29(3), 479–484. 10.1016/j.nut.2012.07.002

Chan, S. Y., Choo, W. S., Young, D. J., & Loh, X. J. (2017). Pectin as a rheology modifier: Origin, structure, commercial production and rheology. Carbohydrate Polymers, 161, 118–139. 10.1016/j.carbpol.2016.12.033

Chang, J., Guo, Y., Yan, J., Zhang, Z., Yuan, L., Wei, C., Zhang, Y., Ma, J., Yang, J., Zhang, X., & Li, H. (2021). The role of watermelon caffeic acid O-methyltransferase (ClCOMT1) in melatonin biosynthesis and abiotic stress tolerance. Horticulture Research, 8(1), 1–12. 10.1038/s41438-021-00645-5

Cohen, R., Burger, Y., Horev, C., Porat, A., & Edelstein, M. (2005). Performance of Galia-type melons grafted on to Cucurbita rootstock in *Monosporascus cannonballus*-infested and non-infested soils. Annals of Applied Biology, 146(3), 381–387. 10.1111/j.1744-7348.2005.040010.x

Cohen, R., Pivonia, S., Crosby, K. M., & Martyn, R. D. (2012). Advances in the biology and management of Monosporascus vine decline and wilt of melons and other cucurbits. In Horticultural Reviews (pp. 77–120). John Wiley & Sons, Ltd. 10.1002/9781118100592.ch2

de Cara, M., López, V., Córdoba, M. C., Santos, M., Jordá, C., & Tello, J. C. (2008). Association of Olpidium bornovanus and Melon necrotic spot virus with Vine Decline of Melon in Guatemala. Plant Disease, 92(5), 709–713. 10.1094/PDIS-92-5-0709

El-Bassossy, H. M., El-Fawal, R., & Fahmy, A. (2012). Arginase inhibition alleviates hypertension associated with diabetes: Effect on endothelial dependent relaxation and NO production. Vascular Pharmacology, 57(5), 194–200. 10.1016/j.vph.2012.01.001

Fenn, M. A., & Giovannoni, J. J. (2021). Phytohormones in fruit development and maturation. The Plant Journal, 105(2), 446–458. 10.1111/tpj.15112

Fish, W. W. (2012). A reliable methodology for quantitative extraction of fruit and vegetable physiological amino acids and their subsequent analysis with commonly available HPLC systems. Food and Nutrition Sciences, 3(6), 863– 871. 10.4236/fns.2012.36115

Fu, L., Xu, B.-T., Xu, X.-R., Gan, R.-Y., Zhang, Y., Xia, E.-Q., & Li, H.-B. (2011). Antioxidant capacities and total phenolic contents of 62 fruits. Food Chemistry, 129(2), 345–350. 10.1016/j.foodchem.2011.04.079

Fulgoni, K., & Fulgoni, V. L. (2022). Watermelon intake is associated with increased nutrient intake and higher diet quality in adults and children, NHANES 2003– 2018. Nutrients, 14(22), 4883. 10.3390/nu14224883

Ghosh, U. K., Islam, M. N., Siddiqui, M. N., Cao, X., & Khan, M. A. R. (2022). Proline, a multifaceted signalling molecule in plant responses to abiotic stress: Understanding the physiological mechanisms. Plant Biology, 24(2), 227–239. 10.1111/plb.13363

Glenn, J. M., Gray, M., Wethington, L. N., Stone, M. S., Stewart, R. W., & Moyen, N. E. (2017). Acute citrulline malate supplementation improves upper- and lower-body submaximal weightlifting exercise performance in resistance-trained females. European Journal of Nutrition, 56(2), 775–784. 10.1007/s00394-015-1124-6

Grabowska, M., Wawrzyniak, D., Rolle, K., Chomczyński, P., Oziewicz, S., Jurga, S., & Barciszewski, J. (2019). Let food be your medicine: Nutraceutical properties of lycopene. Food & Function, 10(6), 3090–3102. 10.1039/C9FO00580C

Hartman, J. L., Wehner, T. C., Ma, G., & Perkins-Veazie, P. (2019). Citrulline and arginine content of taxa of Cucurbitaceae. Horticulturae, 5(1), 22. 10.3390/horticulturae5010022

Haytovitz, D. B., Wu, X., & Bhagwat, S. (2018). *Haytowitz: USDA database for the flavonoid content*. https://agdatacommons.nal.usda.gov/articles/dataset/USDA_Database_for_the_Flavonoid_Content_of_Selected_Foods_Release_3_1_May_2014_/24659802/3

Huang, J., Zou, T., Hu, H., Xiao, X., Wang, Z., Li, M., & Dai, S. (2022). Automatic Brix measurement for watermelon breeding. Applied Sciences, 12(23), 12227. 10.3390/app122312227

Huang, W., Shi, Y., Yan, H., Wang, H., Wu, D., Grierson, D., & Chen, K. (2023). The calcium-mediated homogalacturonan pectin complexation in cell walls contributes the firmness increase in loquat fruit during postharvest storage. Journal of Advanced Research, 49, 47–62. 10.1016/j.jare.2022.09.009

Ilahy, R., Tlili, I., Siddiqui, M. W., Hdider, C., & Lenucci, M. S. (2019). Inside and beyond color: Comparative overview of functional quality of tomato and watermelon fruits. Frontiers in Plant Science, 10, 769. 10.3389/fpls.2019.00769

Islam, J., Shirakawa, H., Nguyen, T. K., Aso, H., & Komai, M. (2016). Simultaneous analysis of serotonin, tryptophan and tryptamine levels in common fresh fruits and vegetables in Japan using fluorescence HPLC. Food Bioscience, 13, 56–59. 10.1016/j.fbio.2015.12.006

Itam, M., Mega, R., Tadano, S., Abdelrahman, M., Matsunaga, S., Yamasaki, Y., Akashi, K., & Tsujimoto, H. (2020). Metabolic and physiological responses to progressive drought stress in bread wheat. Scientific Reports, 10(1), 17189. 10.1038/s41598-020-74303-6

Jang, Y., Huh, Y.-C., Park, D.-K., Mun, B., Lee, S., & Um, Y. (2014). Greenhouse evaluation of melon rootstock resistance to *Monosporascus* root rot and vine decline as well as of yield and fruit quality in grafted “Inodorus” melons. Horticultural Science & Technology, 32(5), 614–622. 10.7235/hort.2014.14065

Kawasaki, S., Miyake, C., Kohchi, T., Fujii, S., Uchida, M., & Yokota, A. (2000). Responses of wild watermelon to drought stress: Accumulation of an ArgE homologue and citrulline in leaves during water deficits. Plant and Cell Physiology, 41(7), 864–873. 10.1093/pcp/pcd005

Levi, A., Jarret, R., Kousik, S., Patrick Wechter, W., Nimmakayala, P., & Reddy, U. K. (2017). Genetic resources of watermelon. In R. Grumet, N. Katzir, & J. Garcia-Mas (Eds.), Genetics and Genomics of Cucurbitaceae (pp. 87–110). Springer International Publishing. 10.1007/7397_2016_34

Liu, J., Guo, S., He, H., Zhang, H., Gong, G., Ren, Y., & Xu, Y. (2013). Dynamic characteristics of sugar accumulation and related enzyme activities in sweet and non-sweet watermelon fruits. Acta Physiologiae Plantarum, 35(11), 3213–3222. 10.1007/s11738-013-1356-0

Luca, S. V., Macovei, I., Bujor, A., Miron, A., Skalicka-Woźniak, K., Aprotosoaie, A. C., & Trifan, A. (2020). Bioactivity of dietary polyphenols: The role of metabolites. Critical Reviews in Food Science and Nutrition, 60(4), 626–659. 10.1080/10408398.2018.1546669

Ludwig, R. A. (1993). Arabidopsis chloroplasts dissimilate L-arginine and L-citrulline for use as N source. Plant Physiology, 101(2), 429–434. 10.1104/pp.101.2.429

Maalej, A., Bouallagui, Z., Hadrich, F., Isoda, H., & Sayadi, S. (2017). Assessment of *Olea europaea* L. fruit extracts: Phytochemical characterization and anticancer pathway investigation. Biomedicine & Pharmacotherapy = Biomedecine & Pharmacotherapie, 90, 179–186. 10.1016/j.biopha.2017.03.034

Maeda, H., & Dudareva, N. (2012). The shikimate pathway and aromatic amino acid biosynthesis in plants. Annual Review of Plant Biology, 63, 73–105. 10.1146/annurev-arplant-042811-105439

Mandal, M. K., Suren, H., Ward, B., Boroujerdi, A., & Kousik, C. (2018). Differential roles of melatonin in plant-host resistance and pathogen suppression in cucurbits. Journal of Pineal Research, 65(3), e12505. 10.1111/jpi.12505

Martyn, R. D. (1996). Monosporascus root rot and vine decline: An emerging disease of melons worldwide. Plant Disease, 80(7), 716. 10.1094/PD-80-0716

Mattioli, R., Costantino, P., & Trovato, M. (2009). Proline accumulation in plants. Plant Signaling & Behavior, 4(11), 1016–1018. 10.4161/psb.4.11.9797

Morita, M., Sakurada, M., Watanabe, F., Yamasaki, T., Doi, H., Ezaki, H., Morishita, K., & Miyake, T. (2013). Effects of oral L-citrulline supplementation on lipoprotein oxidation and endothelial dysfunction in humans with vasospastic angina. Immunology, Endocrine & Metabolic Agents in Medicinal Chemistry, 13(3), 214–220. 10.2174%2F18715222113139990008

Mushtaq, M., Sultana, B., Bhatti, H. N., & Asghar, M. (2015). RSM based optimized enzyme-assisted extraction of antioxidant phenolics from underutilized watermelon (*Citrullus lanatus* Thunb.) rind. Journal of Food Science and Technology, 52(8), 5048–5056. 10.1007/s13197-014-1562-9

Perkins-Veazie, P., Collins, J. K., Pair, S. D., & Roberts, W. (2001). Lycopene content differs among red-fleshed watermelon cultivars. Journal of the Science of Food and Agriculture, 81(10), 983–987. 10.1002/jsfa.880

Pivonia, S., Cohen, R., Katan, J., & Kigel, J. (2002). Effect of fruit load on the water balance of melon plants infected with *Monosporascus cannonballus*. Physiological and Molecular Plant Pathology, 60(1), 39–49. 10.1006/pmpp.2001.0375

R Core Team. (2016). R: A language and environment for statistical computing. [R Foundation for Statistical Computing, Vienna, Austria.]. https://www.R-project.org/

Ren, Y., Liao, S., & Xu, Y. (2023). An update on sugar allocation and accumulation in fruits. Plant Physiology, 193(2), 888–899. 10.1093/plphys/kiad294

Rhouma, A., Salem, I. B., M’hamdi, M., & Boughalleb-M’Hamdi, N. (2019). Relationship study among soils physico-chemical properties and *Monosporascus cannonballus* ascospores densities for cucurbit fields in Tunisia. European Journal of Plant Pathology, 153(1), 65–78. 10.1007/s10658-018-1541-5

Sharma, O. P., & Bhat, T. K. (2009). DPPH antioxidant assay revisited. Food Chemistry, 113(4), 1202–1205. 10.1016/j.foodchem.2008.08.008

Song, Q., Joshi, M., DiPiazza, J., & Joshi, V. (2020). Functional relevance of citrulline in the vegetative tissues of watermelon during abiotic stresses. Frontiers in Plant Science, 11, 512. 10.3389/fpls.2020.00512

Sorokina, M., McCaffrey, K. S., Deaton, E. E., Ma, G., Ordovás, J. M., Perkins-Veazie, P. M., Steinbeck, C., Levi, A., & Parnell, L. D. (2021). A catalog of natural products occurring in watermelon—*Citrullus lanatus*. Frontiers in Nutrition, 8, 729822. 10.3389/fnut.2021.729822

Stanghellini, M. E., Mathews, D. M., & Misaghi, I. J. (2010). Pathogenicity and Management of Olpidium bornovanus, a Root Pathogen of Melons. Plant Disease, 94(2), 163–166. 10.1094/PDIS-94-2-0163

Stanghellini, M. E., Mohammadi, M., & Adaskaveg, J. E. (2014). Effect of soil matric water potentials on germination of ascospores of Monosporascus cannonballus and colonization of melon roots by zoospores of Olpidium bornovanus. European Journal of Plant Pathology, 139(2), 393–398. 10.1007/s10658-014-0395-8

Sun, T., Rao, S., Zhou, X., & Li, L. (2022). Plant carotenoids: Recent advances and future perspectives. Molecular Horticulture, 2(1), 3. 10.1186/s43897-022-00023-2

Sun, T., Yuan, H., Cao, H., Yazdani, M., Tadmor, Y., & Li, L. (2018). Carotenoid metabolism in plants: The role of plastids. Molecular Plant, 11(1), 58–74. 10.1016/j.molp.2017.09.010

Szabados, L., & Savouré, A. (2010). Proline: A multifunctional amino acid. Trends in Plant Science, 15(2), 89–97. 10.1016/j.tplants.2009.11.009

Tarazona-Díaz, M. P., Alacid, F., Carrasco, M., Martínez, I., & Aguayo, E. (2013). Watermelon juice: Potential functional drink for sore muscle relief in athletes. Journal of Agricultural and Food Chemistry, 61(31), 7522–7528. 10.1021/jf400964r

Trovato, M., Funck, D., Forlani, G., Okumoto, S., & Amir, R. (2021). Editorial: Amino acids in plants: Regulation and functions in development and stress defense. Frontiers in Plant Science, 12, 772810. 10.3389/fpls.2021.772810

Tsuboi, T., Maeda, M., & Hayashi, T. (2018). Administration of L-arginine plus L-citrulline or L-citrulline alone successfully retarded endothelial senescence. PLoS ONE, 13(2), e0192252. 10.1371/journal.pone.0192252

Valenta, K., Kalbitzer, U., Razafimandimby, D., Omeja, P., Ayasse, M., Chapman, C. A., & Nevo, O. (2018). The evolution of fruit colour: Phylogeny, abiotic factors and the role of mutualists. Scientific Reports, 8(1), 14302. 10.1038/s41598-018-32604-x

Verbruggen, N., & Hermans, C. (2008). Proline accumulation in plants: A review. Amino Acids, 35(4), 753–759. 10.1007/s00726-008-0061-6

Wang, S.-Y., Shi, X.-C., Wang, R., Wang, H.-L., Liu, F., & Laborda, P. (2020). Melatonin in fruit production and postharvest preservation: A review. Food Chemistry, 320, 126642. 10.1016/j.foodchem.2020.126642

Winkler, A., Fiedler, B., & Knoche, M. (2020). Calcium physiology of sweet cherry fruits. Trees, 34(5), 1157–1167. 10.1007/s00468-020-01986-9

Wong, A., Chernykh, O., & Figueroa, A. (2016). Chronic L-citrulline supplementation improves cardiac sympathovagal balance in obese postmenopausal women: A preliminary report. Autonomic Neuroscience: Basic and Clinical, 198, 50–53. 10.1016/j.autneu.2016.06.005

Wu, X., Ren, J., Huang, X., Zheng, X., Tian, Y., Shi, L., Dong, P., & Li, Z. (2021). Melatonin: Biosynthesis, content, and function in horticultural plants and potential application. Scientia Horticulturae, 288, 110392. 10.1016/j.scienta.2021.110392

Yamada, M., Malambane, G., Yamada, S., Suharsono, S., Tsujimoto, H., Moseki, B., & Akashi, K. (2018). Differential physiological responses and tolerance to potentially toxic elements in biodiesel tree *Jatropha curcas*. Scientific Reports, 8(1), 1635. 10.1038/s41598-018-20188-5

Yokoyama, R., de Oliveira, M. V. V., Kleven, B., & Maeda, H. A. (2021). The entry reaction of the plant shikimate pathway is subjected to highly complex metabolite-mediated regulation. The Plant Cell, 33(3), 671–696. 10.1093/plcell/koaa042

Yuan, P., Umer, M. J., He, N., Zhao, S., Lu, X., Zhu, H., Gong, C., Diao, W., Gebremeskel, H., Kuang, H., & Liu, W. (2021). Transcriptome regulation of carotenoids in five flesh-colored watermelons (*Citrullus lanatus*). BMC Plant Biology, 21(1), 203. 10.1186/s12870-021-02965-z

Zhang, J., Sun, H., Guo, S., Ren, Y., Li, M., Wang, J., Zhang, H., Gong, G., & Xu, Y. (2020). Decreased protein abundance of lycopene β-cyclase contributes to red flesh in domesticated watermelon. Plant Physiology, 183(3), 1171–1183. 10.1104/pp.19.01409

Zheng, Y., Cabassa-Hourton, C., Planchais, S., Lebreton, S., & Savouré, A. (2021). The proline cycle as an eukaryotic redox valve. Journal of Experimental Botany, 72(20), 6856–6866. 10.1093/jxb/erab361

