## Supplementary Information for "Influence of vine decline disease on the amino acid metabolism of watermelon fruit"

**Supplementary Document S1.**

**LC-MS/MS condition**

| | |
|---|---|
| Column temperature: | 40°C |
| Flow speed: | 250 μl $min^{-1}$ |
| Injection volume: | 3 μl |
| Scan Type: | Dynamic MRM |
| Ion source: | ESI |
| Cycle time: | 500 ms |
| MRM repeats: | 3 |

Solvent composition:

| | |
|---|---|
| Solution-A: | 100% water |
| Solvent-B: | 100% acetonitrile |

Gradient timetable

| Step | Time | A | B |
|---|---|---|---|
| 1 | 2 min | 100% | 0% |
| 2 | 5 min | 75% | 25% |
| 3 | 11 min | 65% | 35% |
| 4 | 15 min | 5% | 95% |
| 5 | 18 min | 5% | 95% |

**Supplementary Table S1. Reaction monitoring conditions in LC-MS/MS analysis**

| Compound | Abbr. | Formula | Mw. | Solvent | Polarity | Precursor ion | Frag (V) | Prod ion 1 (CE)† | Prod ion 2 (CE) † | Prod ion 3 (CE) † | Prod ion 4 (CE) † | Ret time (min) |
|---|---|---|---|---|---|---|---|---|---|---|---|---|
| Taurine | Tau | C2H7NO3S | 125.01 | 0.1% formic acid in water | Positive | 126 | 110/90 | 108(9) | 44.2(21) | | | 2.79 |
| Urea | Urea | CH4N2O | 60.06 | 0.1% formic acid in water | Positive | 61 | 50 | 44.2(21) | | | | 3.01 |
| Cystine | Cys-Cys | C6H12N2O4S2 | 240.29 | 0.1% formic acid in water | Positive | 241 | 90 | 151.9(9) | 119.9(17) | 74.1(29) | | 3.11 |
| Allantoin | Alla | C4H6N4O3 | 158.04 | 0.1% formic acid in water | Positive | 159 | 80 | 116.1(5) | 61.2(5) | 44.2(55) | | 3.16 |
| Asparagine | Asn | C4H8N2O3 | 132.12 | 0.1% formic acid in water | Positive | 133.1 | 80 | 87.1(5) | 74.1(13) | 28.3(29) | | 3.2 |
| Serine | Ser | C3H7NO3 | 105.09 | 0.1% formic acid in water | Positive | 106.1 | 70 | 88.1(5) | 60.2(9) | 42.2(25) | | 3.21 |
| Aspartic acid | Asp | C4H7NO4 | 133.11 | 0.1% formic acid in water | Positive | 134 | 70 | 88.1(5) | 74.1(13) | 43.2(25) | | 3.22 |
| Hydroxy-L-proline | Hyp | C5H9NO3 | 131.13 | 0.1% formic acid in 50% MeOH | Positive | 132.1 | 90 | 86.1(13) | 68.2(21) | 41.2(33) | 39.2(55) | 3.3 |
| Glycine | Gly | C2H5NO2 | 75.07 | 0.1% formic acid in water | Positive | 76 | 20 | 48.2(5) | 30.3(5) | | | 3.31 |
| Glutamine | Gln | C5H10N2O3 | 146.14 | 0.1% formic acid in water | Positive | 147.1 | 70 | 130(5) | 84.1(17) | 56.2(33) | | 3.4 |
| Cysteine | Cys | C3H7NO2S | 121.16 | 0.1% formic acid in water | Positive | 122 | 60 | 76.1(13) | 59.1(25) | 43.2(33) | | 3.52 |
| Threonine | Thr | C4H9NO3 | 119.12 | 0.1% formic acid in water | Positive | 120.1 | 70 | 102(5) | 74.1(9) | 56.2(17) | | 3.54 |
| Methionine Sulfoxide | MetSO | C5H11NO3S | 165.05 | 0.1% formic acid in water | Positive | 166.1 | 80 | 75.1(5) | 74.1(13) | 56.2(25) | | 3.63 |
| Alanine | Ala | C3H7NO2 | 89.09 | 0.1% formic acid in water | Positive | 90.1 | 40 | 45.1(41) | 44.2(9) | 29.2(45) | | 3.74 |
| Glutamic acid | Glu | C5H9NO4 | 147.13 | 0.1% formic acid in water | Positive | 148.1 | 80 | 130(5) | 84.1(17) | 56.2(33) | | 3.78 |
| Betaine | Bet | C5H11NO2 | 117.08 | 0.1% formic acid in water | Positive | 118.1 | 120 | 59.2(17) | 58.2(33) | 42.2(55) | | 3.81 |
| Citrulline | Cit | C6H13N3O3 | 175.1 | 0.1% formic acid in water | Positive | 176.1 | 80 | 159.0(5) | 113.0(13) | 70.2(25) | | 3.9 |
| Proline | Pro | C5H9NO2 | 115.13 | 0.1% formic acid in water | Positive | 116.1 | 90 | 70.2(17) | 43.2(37) | 28.3(41) | | 4.27 |
| Ornithine | Orn | C5H12N2O2 | 132.09 | 0.1% formic acid in water | Positive | 133.1 | 80 | 116.0(5) | 70.2(17) | 43.2(37) | | 4.9 |
| beta-Alanine | bAla | C3H7NO2 | 89.05 | 0.1% formic acid in water | Positive | 90.1 | 50 | 72.2(5) | 45.2(45) | 30.3(9) | | 5.1 |
| Histidine | His | C6H9N3O2 | 155.15 | 0.1% formic acid in water | Positive | 156.1 | 100 | 110(13) | 93.1(25) | 83.1(29) | | 5.11 |
| Lysine | Lys | C6H14N2O2 | 146.19 | 0.1% formic acid in water | Positive | 147.1 | 80 | 130(5) | 84.2(17) | 56.2(33) | | 5.2 |
| Arginine | Arg | C6H14N4O2 | 174.2 | 0.1% formic acid in water | Positive | 175.1 | 100 | 116(13) | 70.2(25) | 60.2(13) | | 6.11 |
| Gamma-aminobutyric acid | GABA | C4H9NO2 | 103.12 | 0.1% formic acid in water | Positive | 104.1 | 70 | 87.1(9) | 45.2(25) | 43.2(17) | | 6.54 |
| Cholin | Cho | C5H14NO | 104.11 | 0.1% formic acid in water | Positive | 105.1 | 110 | 46.2(25) | 45.2(21) | 44.2(41) | | 7.34 |
| Putrescine | Put | C4H12N2 | 88.1 | 0.1% formic acid in water | Positive | 89.1 | 70 | 72.2(5) | | | | 7.5 |
| Valine | Val | C5H11NO2 | 117.15 | 0.1% formic acid in water | Positive | 118.1 | 70 | 72.2(9) | 55.2(21) | 42.2(49) | | 7.65 |
| Methionine | Met | C5H11NO2S | 149.21 | 0.1% formic acid in water | Positive | 150.1 | 80 | 133(5) | 104(9) | 56.2(17) | | 8.02 |
| Tyrosine | Tyr | C9H11NO3 | 181.19 | 0.1% formic acid in water | Positive | 182.1 | 80 | 165(5) | 136(9) | 91.1(33) | | 8.83 |
| Isoleucine | Ile | C6H13NO2 | 131.17 | 0.1% formic acid in water | Positive | 132.1 | 80 | 86.2(9) | 44.2(25) | 41.2(29) | | 9.91 |
| Leucine | Leu | C6H13NO2 | 131.17 | 0.1% formic acid in water | Positive | 132.1 | 70 | 86.2(5) | 44.2(25) | 43.2(25) | 30.3(17) | 10.4 |
| Phenylalanine | Phe | C9H11NO2 | 165.19 | 0.1% formic acid in water | Positive | 166.1 | 80 | 120(9) | 103(29) | 77.1(45) | | 11.13 |
| Tryptophan | Trp | C11H12N2O2 | 204.23 | 0.1% formic acid in water | Positive | 205.1 | 90 | 118(29) | 146(13) | 188(5) | | 14.91 |

† Product ion and collision energy

**Supplementary Table S2. Accumulation level of free amino acids in the fruits (mg $gFW^{-1}$)**

| Condition | ID/stat[†] | Cit | Gln | Arg | Glu | Asp | Ile | Val | Ala | Phe | GABA | bAla | Ser | MetO | His | Leu | Met | Thr | Cho | Asn | Tyr | Pro | Trp | Total |
|---|---|---|---|---|---|---|---|---|---|---|---|---|---|---|---|---|---|---|---|---|---|---|---|---|
| Control | ct1 | 8.069 | 4.927 | 3.801 | 0.853 | 1.050 | 1.099 | 1.103 | 0.874 | 0.929 | 0.865 | 0.710 | 0.492 | 0.402 | 0.472 | 0.342 | 0.324 | 0.193 | 0.173 | 0.133 | 0.078 | 0.089 | 0.093 | 27.072 |
| | ct2 | 7.516 | 5.333 | 4.454 | 1.685 | 1.077 | 1.229 | 1.174 | 0.946 | 0.899 | 0.903 | 0.887 | 0.456 | 0.399 | 0.363 | 0.372 | 0.289 | 0.218 | 0.187 | 0.089 | 0.076 | 0.067 | 0.058 | 28.678 |
| | ct3 | 6.476 | 4.747 | 3.632 | 1.538 | 1.212 | 0.996 | 1.047 | 0.986 | 0.760 | 0.804 | 0.829 | 0.410 | 0.386 | 0.275 | 0.294 | 0.265 | 0.204 | 0.145 | 0.105 | 0.064 | 0.102 | 0.049 | 25.327 |
| | ct4 | 6.698 | 4.290 | 3.736 | 1.352 | 0.877 | 1.004 | 0.972 | 0.777 | 0.744 | 0.746 | 0.718 | 0.297 | 0.378 | 0.263 | 0.321 | 0.188 | 0.172 | 0.160 | 0.061 | 0.071 | 0.028 | 0.033 | 23.886 |
| | ct5 | 6.624 | 4.658 | 3.611 | 1.491 | 1.209 | 1.047 | 1.058 | 0.791 | 0.867 | 0.712 | 0.786 | 0.391 | 0.283 | 0.353 | 0.324 | 0.317 | 0.185 | 0.123 | 0.082 | 0.078 | 0.086 | 0.094 | 25.168 |
| | mean | 7.076 | 4.791 | 3.847 | 1.384 | 1.085 | 1.075 | 1.071 | 0.875 | 0.840 | 0.806 | 0.786 | 0.409 | 0.369 | 0.345 | 0.331 | 0.277 | 0.194 | 0.157 | 0.094 | 0.074 | 0.074 | 0.065 | 26.026 |
| | SD[‡] | 0.614 | 0.341 | 0.311 | 0.286 | 0.123 | 0.085 | 0.067 | 0.082 | 0.075 | 0.071 | 0.067 | 0.066 | 0.044 | 0.075 | 0.026 | 0.049 | 0.016 | 0.022 | 0.024 | 0.005 | 0.026 | 0.024 | 1.669 |
| MRRVD | vd1 | 5.653 | 1.116 | 2.404 | 0.891 | 0.369 | 0.795 | 0.786 | 0.700 | 0.758 | 0.324 | 0.427 | 0.173 | 0.000 | 0.243 | 0.213 | 0.212 | 0.072 | 0.068 | 0.000 | 0.029 | 0.077 | 0.115 | 15.426 |
| | vd2 | 7.432 | 2.651 | 2.810 | 1.307 | 0.943 | 1.027 | 0.936 | 0.408 | 0.955 | 0.485 | 0.653 | 0.259 | 0.241 | 0.330 | 0.259 | 0.306 | 0.165 | 0.100 | 0.074 | 0.045 | 0.110 | 0.109 | 21.606 |
| | vd3 | 8.432 | 1.184 | 2.724 | 0.563 | 0.675 | 0.994 | 0.986 | 0.438 | 1.044 | 0.572 | 0.416 | 0.163 | 0.254 | 0.387 | 0.253 | 0.226 | 0.119 | 0.140 | 0.065 | 0.037 | 0.215 | 0.088 | 19.973 |
| | vd4 | 8.602 | 2.316 | 3.066 | 1.042 | 1.050 | 0.972 | 0.924 | 0.265 | 1.002 | 0.825 | 0.633 | 0.201 | 0.247 | 0.366 | 0.262 | 0.286 | 0.203 | 0.121 | 0.078 | 0.060 | 0.203 | 0.110 | 22.836 |
| | vd5 | 8.602 | 2.425 | 2.491 | 0.811 | 0.788 | 0.915 | 0.876 | 0.250 | 0.919 | 0.586 | 0.610 | 0.252 | 0.355 | 0.429 | 0.224 | 0.267 | 0.276 | 0.157 | 0.080 | 0.044 | 0.197 | 0.072 | 21.627 |
| | mean | 7.744 | 1.939 | 2.699 | 0.923 | 0.765 | 0.940 | 0.902 | 0.412 | 0.936 | 0.558 | 0.548 | 0.210 | 0.219 | 0.351 | 0.242 | 0.259 | 0.167 | 0.117 | 0.059 | 0.043 | 0.160 | 0.099 | 20.293 |
| | SD[‡] | 1.133 | 0.653 | 0.236 | 0.247 | 0.236 | 0.081 | 0.068 | 0.162 | 0.098 | 0.163 | 0.104 | 0.040 | 0.117 | 0.063 | 0.020 | 0.036 | 0.070 | 0.031 | 0.030 | 0.010 | 0.056 | 0.016 | 2.598 |
| P-value[¶] | | 0.330 | **0.000** | **0.000** | **0.041** | **0.043** | 0.052 | **0.007** | **0.001** | 0.159 | **0.024** | **0.005** | **0.001** | **0.044** | 0.910 | **0.001** | 0.584 | 0.468 | 0.068 | 0.110 | **0.001** | **0.024** | 0.051 | **0.006** |

[†] Plant IDs are shown by two alphabet letters (ct or vd for control and MRRVD-affected watermelon) followed by one digit. Statistical terms are also indicated.

[‡] Standard deviation.

[¶] Probability value by Student's *t*-test between control and MRRVD groups. Values less than 0.05 are indicated in bold.

**Supplementary Table S3. Percentage fraction of free amino acids in the fruits (%)**

| Condition | ID/stat[*1] | Cit | Gln | Arg | Glu | Asp | Ile | Val | Ala | Phe | GABA | bAla | Ser | MetO | His | Leu | Met | Thr | Cho | Asn | Tyr | Pro | Trp |
|---|---|---|---|---|---|---|---|---|---|---|---|---|---|---|---|---|---|---|---|---|---|---|---|
| Control | ct1 | 29.81 | 18.20 | 14.04 | 3.15 | 3.88 | 4.06 | 4.08 | 3.23 | 3.43 | 3.20 | 2.62 | 1.82 | 1.48 | 1.74 | 1.26 | 1.20 | 0.71 | 0.64 | 0.49 | 0.29 | 0.33 | 0.34 |
| | ct2 | 26.21 | 18.59 | 15.53 | 5.88 | 3.76 | 4.29 | 4.09 | 3.30 | 3.14 | 3.15 | 3.09 | 1.59 | 1.39 | 1.26 | 1.30 | 1.01 | 0.76 | 0.65 | 0.31 | 0.27 | 0.23 | 0.20 |
| | ct3 | 25.57 | 18.74 | 14.34 | 6.07 | 4.78 | 3.93 | 4.14 | 3.89 | 3.00 | 3.18 | 3.27 | 1.62 | 1.52 | 1.09 | 1.16 | 1.05 | 0.81 | 0.57 | 0.42 | 0.25 | 0.40 | 0.19 |
| | ct4 | 28.04 | 17.96 | 15.64 | 5.66 | 3.67 | 4.20 | 4.07 | 3.25 | 3.11 | 3.12 | 3.01 | 1.24 | 1.58 | 1.10 | 1.35 | 0.79 | 0.72 | 0.67 | 0.26 | 0.30 | 0.12 | 0.14 |
| | ct5 | 26.32 | 18.51 | 14.35 | 5.93 | 4.80 | 4.16 | 4.21 | 3.14 | 3.44 | 2.83 | 3.12 | 1.55 | 1.12 | 1.40 | 1.29 | 1.26 | 0.73 | 0.49 | 0.32 | 0.31 | 0.34 | 0.37 |
| | mean | 27.19 | 18.40 | 14.78 | 5.34 | 4.18 | 4.13 | 4.12 | 3.36 | 3.22 | 3.09 | 3.02 | 1.56 | 1.42 | 1.32 | 1.27 | 1.06 | 0.75 | 0.60 | 0.36 | 0.28 | 0.28 | 0.25 |
| | SD[*2] | 1.54 | 0.28 | 0.67 | 1.10 | 0.51 | 0.12 | 0.05 | 0.27 | 0.18 | 0.14 | 0.22 | 0.19 | 0.16 | 0.24 | 0.06 | 0.16 | 0.03 | 0.07 | 0.08 | 0.02 | 0.10 | 0.09 |
| MRRVD | vd1 | 36.64 | 7.24 | 15.58 | 5.78 | 2.39 | 5.15 | 5.09 | 4.54 | 4.92 | 2.10 | 2.77 | 1.12 | 0.00 | 1.58 | 1.38 | 1.37 | 0.46 | 0.44 | 0.00 | 0.19 | 0.50 | 0.75 |
| | vd2 | 34.40 | 12.27 | 13.01 | 6.05 | 4.36 | 4.75 | 4.33 | 1.89 | 4.42 | 2.25 | 3.02 | 1.20 | 1.12 | 1.53 | 1.20 | 1.42 | 0.76 | 0.46 | 0.34 | 0.21 | 0.51 | 0.50 |
| | vd3 | 42.22 | 5.93 | 13.64 | 2.82 | 3.38 | 4.98 | 4.94 | 2.19 | 5.23 | 2.86 | 2.08 | 0.81 | 1.27 | 1.94 | 1.27 | 1.13 | 0.60 | 0.70 | 0.33 | 0.19 | 1.08 | 0.44 |
| | vd4 | 37.67 | 10.14 | 13.43 | 4.56 | 4.60 | 4.26 | 4.05 | 1.16 | 4.39 | 3.61 | 2.77 | 0.88 | 1.08 | 1.60 | 1.15 | 1.25 | 0.89 | 0.53 | 0.34 | 0.26 | 0.89 | 0.48 |
| | vd5 | 39.78 | 11.21 | 11.52 | 3.75 | 3.64 | 4.23 | 4.05 | 1.16 | 4.25 | 2.71 | 2.82 | 1.17 | 1.64 | 1.98 | 1.04 | 1.23 | 1.28 | 0.73 | 0.37 | 0.21 | 0.91 | 0.33 |
| | mean | 38.14 | 9.36 | 13.43 | 4.59 | 3.68 | 4.67 | 4.49 | 2.19 | 4.64 | 2.71 | 2.69 | 1.04 | 1.02 | 1.73 | 1.21 | 1.28 | 0.80 | 0.57 | 0.28 | 0.21 | 0.78 | 0.50 |
| | SD[*2] | 2.67 | 2.40 | 1.31 | 1.22 | 0.78 | 0.37 | 0.44 | 1.24 | 0.37 | 0.53 | 0.32 | 0.16 | 0.55 | 0.19 | 0.12 | 0.10 | 0.28 | 0.12 | 0.14 | 0.03 | 0.23 | 0.14 |
| P-value[*3] | | **0.00010** | **0.00007** | 0.10372 | 0.39016 | 0.31223 | **0.02433** | 0.12940 | 0.10193 | **0.00013** | 0.19710 | 0.12547 | **0.00251** | 0.20028 | **0.03078** | 0.36033 | 0.05041 | 0.72434 | 0.65906 | 0.32863 | **0.00284** | **0.00458** | **0.01549** |

*1: Plant IDs are shown by two alphabet letters (ct or vd for control and MRRVD-affected watermelon) followed by one digit. Statistical terms are also indicated.

*2: Standard deviation.

*3: Probability value by Student's *t*-test between control and MRRVD groups. Values less than 0.05 are indicated in bold.
